# Human cornea harbors tissue-resident memory T cells shaped by systemic immune activation, age and biological sex

**DOI:** 10.64898/2026.07.20.739463

**Authors:** Mengliang Wu, Sarah C. Monard, Marina H. Yakou, Kirthana Senthil, Sapna Devi, Yannick O. Alexandre, Ching Yi Wu, Thomas N. Burn, Joanna R. Groom, Bao N. Nguyen, Laura K. Mackay, Holly R. Chinnery, Scott N. Mueller, Laura E. Downie

**Author notes:** Correspondence to: Laura Downie, Scott Mueller, Holly Chinnery. Joint Senior Authors (Holly R. Chinnery, Scott N. Mueller, Laura E. Downie).

## Abstract

The cornea is classically regarded as an immune-privileged tissue. However, recent studies have identified T cells in the healthy human cornea that are absent in specific pathogen-free (SPF) mice. The identity of these T cells, the systemic cues that drive their establishment, and the molecular mechanisms governing their corneal homing remain unknown. Here, we combine multimodal human data with tractable murine models to characterize the cellular basis of corneal immune surveillance. Immunofluorescence staining and confocal imaging of clinically non-inflamed human donor tissues revealed CD3⁺ T cells within the corneal epithelium. Flow cytometry revealed that these cells were predominantly CD8⁺, with a tissue-resident memory T (T_RM_) cell phenotype. *In vivo* confocal microscopy demonstrated that corneal T cell abundance increased with age, particularly in males, suggesting that accumulation is shaped by cumulative systemic immune experience. Whereas young SPF mice were devoid of corneal T cells, infection with pathogens that do not typically target the cornea induced long-lived T_RM_-like CD8⁺ T cells in the cornea. We identified CXCR6 as required for efficient T cell recruitment to the cornea of viral-infected mice. Together, our human and murine data support a model in which systemic immune history, age and biological sex influence local immune surveillance at the human ocular surface.

## Main

The cornea has long been considered an immune-privileged tissue, where adaptive immune cell traffic is tightly constrained under homeostatic conditions^1^. This view is strongly premised on studies in specific pathogen-free laboratory mice, where the healthy mouse cornea is essentially devoid of T cells in the steady state, and populated instead by myeloid cells^2, 3, 4^. In contrast, we recently revealed the presence of patrolling intraepithelial T cells in the healthy human cornea, which redefines the current conceptualization of the human corneal immune compartment and challenges dogma relating to ocular immune privilege^5, 6^.

With this paradigm-shifting discovery that T cells comprise a major immune cell subset in the steady-state human cornea^7^, there remains a need to address key questions about their immunological identity and to define why the human corneal immune cell niche is distinct from that of mice under homeostatic conditions. It is not known whether human corneal T cells are CD4⁺ or CD8⁺, if they represent tissue-resident populations, and if their abundance or dynamic activity changes across the hosts lifespan. Defining these features is essential to advancing understanding of the immunological landscape of the human cornea, and how homeostatic immune surveillance might relate to susceptibility to ocular disease.

In the murine cornea, local herpes simplex virus (HSV) infection leads to the formation of CD8⁺ tissue-resident memory T (T_RM_) cells that persist long after viral clearance, demonstrating that this tissue can support durable T cell residency following infectious inflammation^8, 9^. This raises important questions of whether T_RM_ cells patrol the human cornea under physiological conditions and, if so, how they are recruited and maintained. At other barrier sites, such as the skin, gut and lung, T_RM_ cells are present in histologically normal tissue^10, 11, 12^, where they provide rapid local protection upon rechallenge. Experimental models further show that systemic viral infections can induce effector CD8^+^ T cells to migrate to distal non-lymphoid organs to form T_RM_ populations, independent of local antigen presence^13, 14^. These findings raise the possibility that cumulative systemic immune experience might populate peripheral tissue sites, including the immune-privileged cornea, with resident T cells, even in the absence of local infection.

To address these questions, we analyzed the phenotype of corneal immune cells in human donor tissues and identified abundant CD8⁺ T_RM_-phenotype cells within the corneal epithelium. Using a novel human corneal intravital imaging technique developed in our laboratory, Functional *In Vivo* Confocal Microscopy (Fun-IVCM)^5, 15^, we discovered sex-dependent differences in the density and motility of human corneal T cells that varied with age. Experiments in SPF mice, which normally lack corneal T_RM_ cells, revealed that systemic and respiratory viral infections can induce long-lived T_RM_ cells in the cornea, via a process dependent on expression of the chemokine receptor CXCR6. Together, this body of work newly identifies an age- and biological sex-dependent compartment of CD8⁺ T_RM_ cells in the human cornea that can be shaped by systemic immune experience. The absence of T_RM_ cells in the steady-state mouse cornea is an important reminder to consider species differences when investigating the function of the immune system in tissue physiology and disease.

## Results

### The healthy human cornea houses populations of T cells

To define the phenotype and distribution of immune cells in human corneas, we first characterized immune cells from post-mortem donor tissues with no known history of ocular infection or trauma, or recent systemic infection (Donor information in Supplemental Table 1). We performed flow cytometric analysis on full thickness human donor corneal tissues (i.e., epithelium and stroma combined) (n=17, 7 females and 10 males, aged 26 to 82 years). The corneal tissues were separated into central and peripheral regions (Supplemental Fig.1a), as previous evidence shows markedly different abundances of immune cells in these regions^16^. To achieve an unbiased view of human corneal immune cell heterogeneity, we performed unsupervised clustering on concatenated flow cytometry data (n=12) and visualized the clusters on a uniform manifold approximation and projection (UMAP) plot. This analysis revealed the presence of CD8^+^ and CD4^+^ T cell subsets, as distinct from the HLA-DR^+^CD14^+^ myeloid lineage (Fig. 1a-b). Further, a population of CD45RA^+^ cells lacking conventional lineage markers was also identified, potentially representing innate lymphoid cells (Fig. 1a-b). Manual gating of immune cell populations confirmed that CD3⁺ T cells were identified in all donor samples (Fig. 1c-e). Within the CD3^-^ immune cell compartment, HLA-DR⁺CD14⁺ myeloid cells were the predominant population (Fig. 1c, f-g), representing macrophages or dendritic cells with a monocyte lineage, with most co-expressing CD11c (Supplemental Fig. 1b-c).

**Figure 1.**
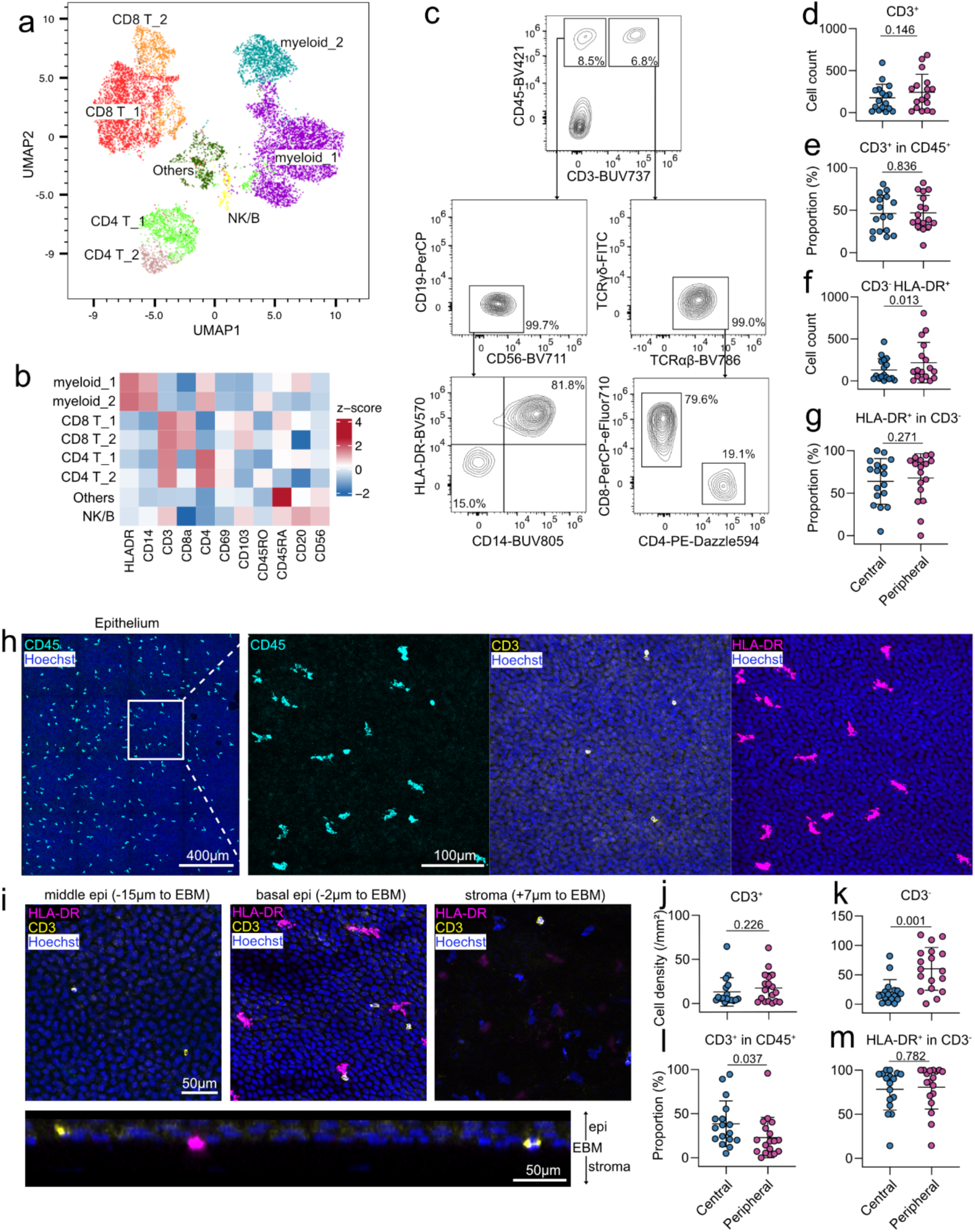
T cell and myeloid cell populations reside in the healthy human corneal epithelium. (a) Unsupervised UMAP clustering analysis on concatenated immune cells from 12 donor corneas. (b) Heatmap showing immune marker levels of different immune cell clusters. (c) Representative flow cytometry plots from an 82-year-old female donor tissue. (d-g) Flow cytometry data comparing central and peripheral corneal regions. More CD3^-^HLA-DR^+^ cells were found in the peripheral cornea relative to the central region. (h) Representative immunofluorescence images of a human cornea from a 31-year-old male donor illustrate a distinct population of CD3^+^ T cells. (i) Representative images from the same donor tissue demonstrate corneal T cells distribution across the epithelium, with myeloid cells mainly residing in the basal epithelium. In the human corneal anterior stroma, myeloid cells dominate the immune compartment, with occasional T cells observed. (j-m) Quantitative analysis plots from immunofluorescence staining data comparing central and peripheral corneal regions. The density of CD3^-^ cells was higher in the peripheral human corneal epithelium than in the central cornea, while the percentage of CD3^+^ cells was higher in the central corneal epithelium. Statistical tests: Paired *t* test (d, e, g, j, k, l and m), paired Wilcoxon test (f). Numbers above graphs indicate P values. Sample numbers: 17 (d-g) and 18 (j-m). Data are shown as mean ± SD. EBM, epithelial basement membrane.

Immunofluorescent staining of corneal flat mounts revealed that CD45⁺ immune cells were sparsely distributed in both the epithelium and stroma (Fig. 1h). Notably, CD3^+^ T cells were identified in the epithelium of all examined corneas (n=18, 8 females and 10 males, aged 29 to 82 years) (Fig. 1h-k), constituting 20-30% of total epithelial immune cells (Fig. 1l). While the density of corneal epithelial T cells showed substantial inter-individual heterogeneity (ranging from 1.4 to 63.7 cells/mm^2^), their cellular distribution was similar across central and peripheral tissue regions (Fig. 1j). In contrast, more CD45⁺CD3⁻ cells were observed in the peripheral, than the central, cornea (Fig. 1k), consistent with evidence from both animal models and clinical reports that myeloid cells have a centripetal gradient and higher densities in the peripheral cornea^16, 17^. The CD3⁻ immune cells were predominantly HLA-DR^+^, suggesting a professional antigen-presenting cell phenotype (Fig. 1h, m). Immune cell evaluation in the corneal stroma was constrained by limited antibody penetration, however abundant CD45⁺ HLA-DR^low^ amoeboid cells (presumed macrophages) and occasional T cells were observed in the anterior stroma (Fig. 1i, Supplemental Fig. 1d). These data establish that the steady-state human cornea houses conventional αβT cells that reside in the basal epithelium.

### Human corneal T cells exhibit tissue-resident memory (T_RM_) phenotypes

We next sought to define whether human corneal T cells represent transient infiltrates or a stable resident cell population. Immunofluorescence imaging revealed that most (∼85%) of the intraepithelial T cells were CD8^+^, while CD4⁺ T cells were less frequent (Fig. 2a-c). A significant proportion of these CD8⁺ T cells co-expressed the canonical residency markers CD103 and CD69 (Fig. 2d-f), a hallmark of T_RM_ cells.

**Figure 2.**
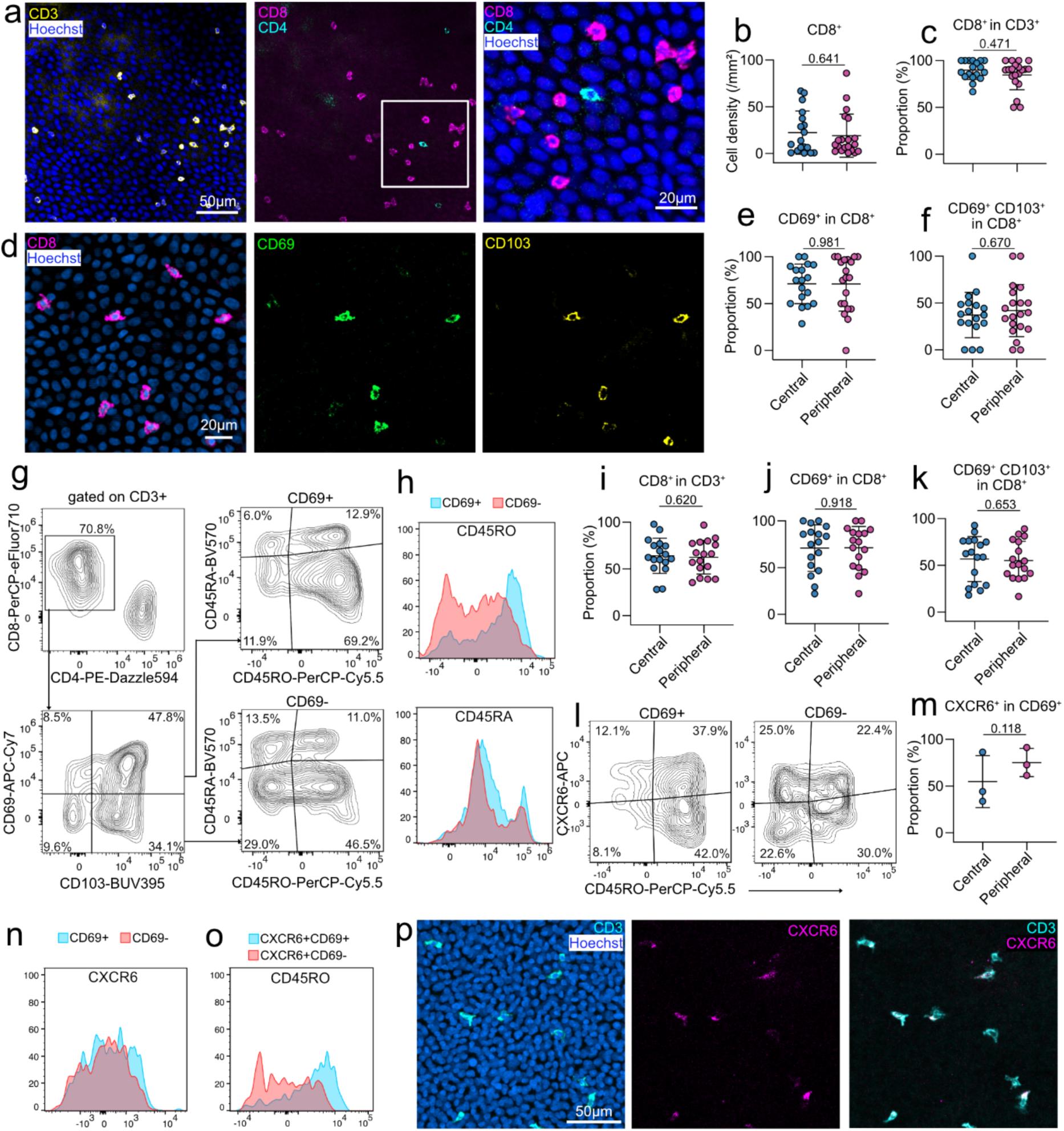
T_RM_ cells populate the healthy human cornea. (a) Representative immunofluorescence images from a 61-year-old male donor showing that CD8^+^ T cells are predominant in the corneal epithelium. (b-c) Quantitative analysis of imaging data showing the CD8^+^ T cell density and their proportion in the central and peripheral human corneal epithelium. (d) Representative immunofluorescence images from a 78-year-old male donor showing CD8^+^CD69^+^CD103^+^ T cells. (e-f) Quantitative analysis of imaging data showing the proportion of CD8^+^CD69^+^CD103^+/-^ T cells in the central and peripheral human corneal epithelium. (g-h) Flow cytometry plots concatenated from 13 donor corneas shows the CD45RA and CD45RO levels in CD69^+^ and CD69^-^ T cell populations. The CD69^+^ population had a higher expression of the memory marker, CD45RO. (i-k) Flow cytometry data reveals the proportion of CD8^+^CD69^+^CD103^+/-^ T cells in combined human corneal epithelium and stroma. (l-o) Flow cytometry analysis concatenated from human donor corneas (n=3) indicates CXCR6 levels in CD69^+^ and CD69^-^ T cell populations and CD45RO levels in CXCR6^+^CD69^+^ and CXCR6^+^CD69^-^ T cell populations. (p) Representative immunofluorescence images from a 78-year-old male donor showing corneal T cells expressing CXCR6. Statistical tests: Paired *t* test (b, e, f, i, j, k, and m), paired Wilcoxon test (c). Numbers above graphs indicate P values. Sample numbers: 20 (b, c, e and f), 17 (i-k) and 3 (m). Data are shown as mean ± SD.

Flow cytometric analysis of full thickness human corneas (n=17) revealed a lower proportion of CD8^+^ T cells (∼60%) compared to the immunostained epithelial samples (Fig. 2g-i), suggesting that CD4^+^ T cells may predominantly reside within the stroma. This compartmentalization mirrors the immune architecture of other barrier tissues, such as the skin and conjunctiva^18, 19^. More than half of the human corneal CD8^+^ T cells co-expressed CD69 and CD103 (Fig. 2j-k), confirming a T_RM_ phenotype. Further analysis showed that the majority of CD69^+^ T cells had a CD45RA⁻CD45RO^+^ memory phenotype (Fig. 2g-h). CD69^+^ T cells expressed higher levels of CD45RO than CD69^−^ cells, consistent with a more memory-like phenotype. Both populations had low CD45RA expression, confirming that the analyzed CD8^+^ T cells were predominantly antigen experienced. In addition, we observed that more than half of the CD4^+^ T cell population expressed CD69 (Supplemental Fig. 1e-f).

Having established that human corneal T cells have a T_RM_-like phenotype, we next sought to investigate whether human corneal T cells express chemokine receptors known to regulate T_RM_ positioning, focusing on CXCR6. The chemokine receptor CXCR6 promotes the long-term survival and retention of T_RM_ cells in barrier tissues like the skin and airways^20, 21^. In the human cornea, we found that CXCR6^+^ subpopulations were equally prevalent (∼50%) among both CD69^+^ and CD69^−^ CD8^+^ T cells (Fig. 2l-n). Within the CXCR6^+^ CD8^+^ T cell pool, the CD69^+^ subset showed higher CD45RO expression (Fig. 2o), further confirming memory maturation in double-positive cells. CXCR6 expression in human corneal T cells was verified by immunofluorescent staining (Fig. 2p). Together, these phenotypic signatures provide evidence for the human cornea housing T_RM_ cell populations, including a dominant population of CD8^+^ T_RM_ cells in the epithelium.

### The human corneal T cell population undergoes age- and sex-dependent expansion

Further to this *ex vivo* analytical profiling of T_RM_ cell populations in human donor corneas, we next sought to characterize the *in vivo* density and temporal dynamics of these cell populations in the corneas of living humans of different sexes and ages. To achieve this, we leveraged a clinical intravital imaging method that we pioneered known as Fun-IVCM that involves the derivation of time-lapsed videos (up to about 25-minutes duration). Quantitative analyses from Fun-IVCM videos defined the *in vivo* distribution and dynamic behaviors of human corneal immune cells, comprising putative motile T cells and sessile dendritic cells (DCs) in the corneal epithelium^5^. Two tissue regions, comprising the whorl (inferonasal, paracentral cornea) and periphery (∼2 mm inferior to the whorl) were imaged in 116 healthy volunteers (60 females and 56 males, aged 18 to 78 years; Fig 3a-b). All participants had no known history of clinically significant ocular infection.

**Figure 3.**
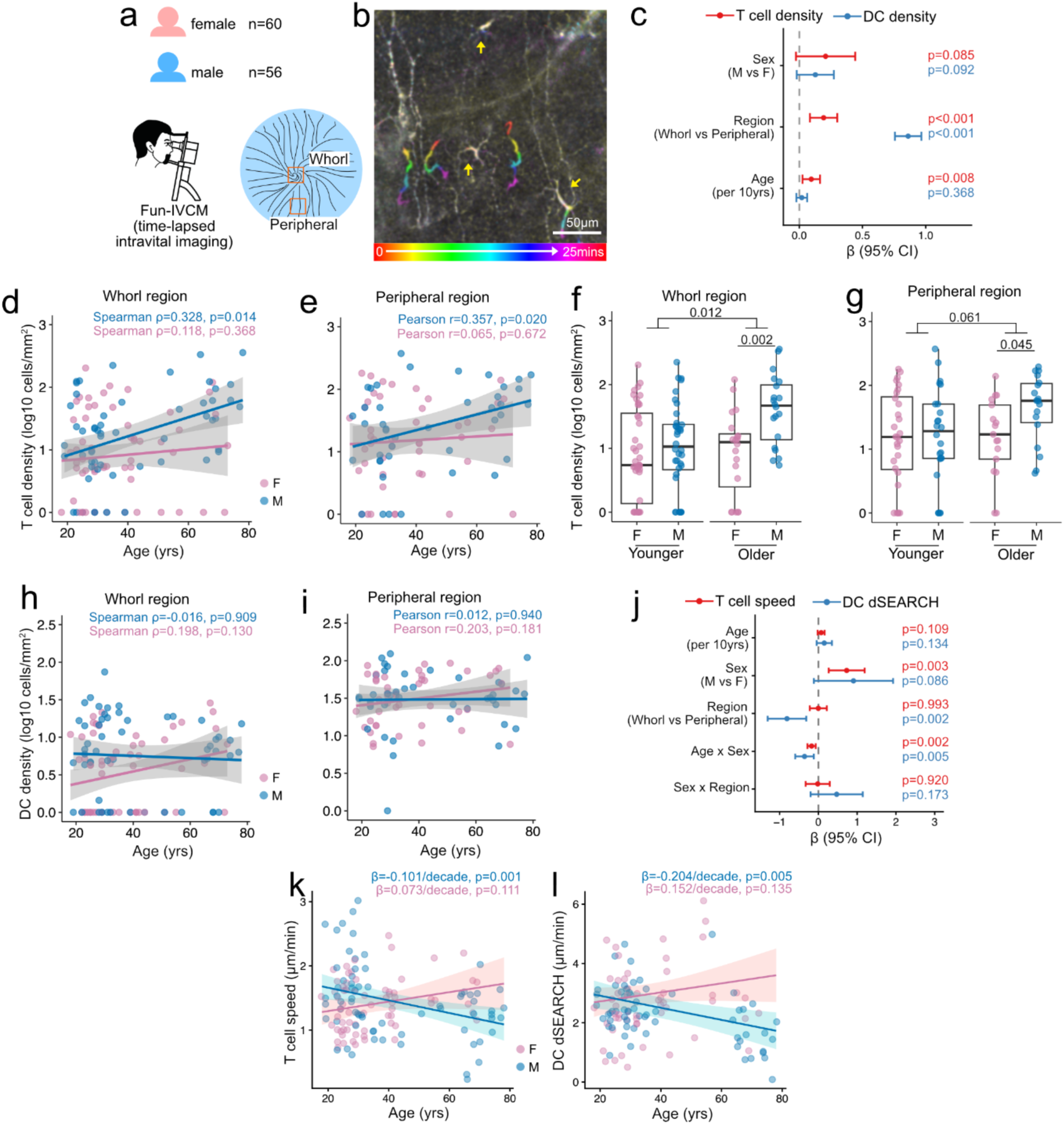
Human corneal T cell populations expand over the lifespan in males. (a) A total of 60 adult females and 56 adult males were included in the Fun-IVCM intravital imaging analysis to derive *in vivo* corneal immune cell data across the lifespan. The whorl (inferonasal paracentral cornea) and peripheral (∼2 mm inferior to the whorl) corneal regions were imaged. (b) Representative time-colored image from a 23-year-old healthy male showing motile T cells (colored tracks) and relatively stationary DCs (arrows) in the human corneal epithelium. (c) Forest plot showing standardized β coefficients (with 95% CIs) from linear mixed-effects models fitted to human corneal *in vivo* immune cell density data for participants of considering the effect of both biological sex and age. (d-e) Correlations between the *in vivo* density of putative human corneal intraepithelial T cells and participant age, stratified by biological sex, in the whorl and peripheral corneal regions. Shaded areas indicate 95% confidence intervals (CIs). (f-g) A higher corneal T cell density was observed in older adults (≥ 40 yrs) compared to younger adults (< 40 yrs); this sex-based difference was observed only in older participants. (h-i) Correlations between the *in vivo* density of putative DCs and participant age, stratified by biological sex, in the whorl and peripheral corneal regions, showing no significant relationship with age. Shaded areas indicate 95% CIs. (j) Forest plot showing standardized β coefficients (with 95% CIs) from linear mixed-effects models fitted to participant-averaged dynamic immune cell data. A significant interaction between biological sex and age was identified for both corneal T cell speed and DC dSEARCH behavior. (k-l) Model-predicted association between participant age and corneal T cell speed or DC dSEARCH, stratified by participant sex. Predicted values were generated while accounting for tissue-region effects. Shaded areas indicate 95% CIs. Statistical tests: Linear mixed-effects model (c, j, k and l), Spearman correlation test (d and h), Pearson correlation test (e and i), Welch’s *t* test (f and g). Numbers above graphs indicate P values. Data (f and g) are shown as median with interquartile range.

The linear mixed-effects model revealed that the *in vivo* density of human corneal epithelial T cells increased with age (β = 0.095, corresponding to a 1.25-fold increase per decade, Fig. 3c; Supplemental Fig. 2a-b), with the peripheral cornea having more T cells than the whorl region (Fig. 3c). Overall, human corneal T cell density trended towards being higher in males than females (Fig. 3c). Per-region tissue analyses confirmed the sex-dependent patterns. In males, a positive correlation between biological age and epithelial T cell density was identified in both the corneal whorl and periphery; this effect was not seen in females (Fig. 3d-e). When participants were dichotomously sub-grouped by age as younger (<40 years) or older (40 years or more), older participants had higher corneal T cell density in the whorl region than younger participants (Fig. 3f-g). A sex-dependent difference was also observed only in older males, whom had more corneal epithelial T cells than females (Fig. 3f-g). In contrast, corneal epithelial DC density showed no association with age nor sex (Fig. 3c, h-i; Supplemental Fig. 2c-f), in addition to a well-documented tissue region effect (Fig. 3c).

We also examined potential age-related effects on the *in vivo* patrolling activity of human corneal T cells by measuring their mean instantaneous speed. At the participant level (T cell speed averaged per participant), there was an interaction effect for age and biological sex in the mixed-effects model (Fig. 3j), with increasing age having a negative effect on cell speed in males (-0.10 µm/min per decade, Fig. 3k) but not in females. No tissue region-dependent differences in T cell speed were observed (Fig. 3j). For corneal DCs, dendrite extension and retraction behavior (dSEARCH index) was measured as an indicator of cell activity. Similarly, a significant interaction between age and sex was identified in the mixed-effects model (Fig. 3j), with increasing age having a negative effect on DC dSEARCH in males (-0.20 µm/min per decade, Fig. 3l) but not in females. Furthermore, an overall tissue region effect was observed, showing a higher DC dSEARCH index in the peripheral cornea than the whorl (Fig. 3j). These findings were supported by an additional analysis using cell-level data in a mixed-effects model (Supplemental Fig.2g)

In summary, we found that advancing age was associated with greater corneal T cell density in people with healthy eyes, which tended to be more apparent in males. In contrast, both human corneal T cell speed and DC dSEARCH, as *in vivo* measures of cell activity, declined with age in males, but remained unchanged in females across the lifespan.

### Systemic immune activation establishes a persistent corneal T_RM_ niche

The positive correlation between age and *in vivo* corneal T cell density in the clinical cohort is consistent with the accumulation of T cells in healthy human corneas over the lifespan, even in individuals without a known history of corneal infection. We have previously shown that local (corneal) infection with Herpes Simplex Virus (HSV) in mice can induce long-lived corneal CD8^+^ T_RM_ cells that can respond *in situ* to antigen rechallenge^8^. Humans are regularly exposed to pathogens and environmental antigens through multiple routes, with systemically and intestinally primed T cells showing the propensity to establish immune memory throughout the body^22, 23^. Therefore, we hypothesized that the observed corneal T_RM_ cell pool might result from systemic immune priming, in addition to being inducted by local infection^8^.

Here, we employed a mouse model of systemic lymphocytic choriomeningitis virus (LCMV, Armstrong strain; Fig. 4a), which induces a self-limited, acute infection that is cleared in 7 to 10 days. At 28 days after acute systemic LCMV infection, we observed an expanded CD8⁺ T cell population in the mouse cornea, including a LCMV-specific P14 cell population with a higher proportion than in spleen (Fig. 4b-f). Most of these P14 cells had an activated effector-memory phenotype (CD44⁺CD62L⁻) (Fig. 4g). Importantly, approximately half of these infiltrating P14 cells co-expressed CD69 and CXCR6, which were nearly absent on splenic P14 cells from the same animals (Fig. 4h). We also noted that CD69^+^ P14 cells in the cornea showed a shift towards a CXCR6⁺PD-1⁺ phenotype, compared with CD69⁻ P14 cells (Fig. 4c). A similar T_RM_ phenotype was also observed amongst endogenous CD8 T cells (Supplemental Fig. 3a). These findings suggest that the corneal environment promotes T_RM_ differentiation by recruited circulating T cells in the absence of local infection. These T cells persisted in the cornea for at least 81 days post-infection (Fig. 4i). The infiltration of T cells into the cornea after systemic LCMV infection was confirmed using immunofluorescence staining at days 12 and 60 (Supplemental Fig. 3b-c). Together, these data identify that systemic viral infection can establish T_RM_ cell populations in the cornea, providing a probable explanation for the presence of T cells in human donors with no known history of ocular injury or infection.

**Figure 4.**
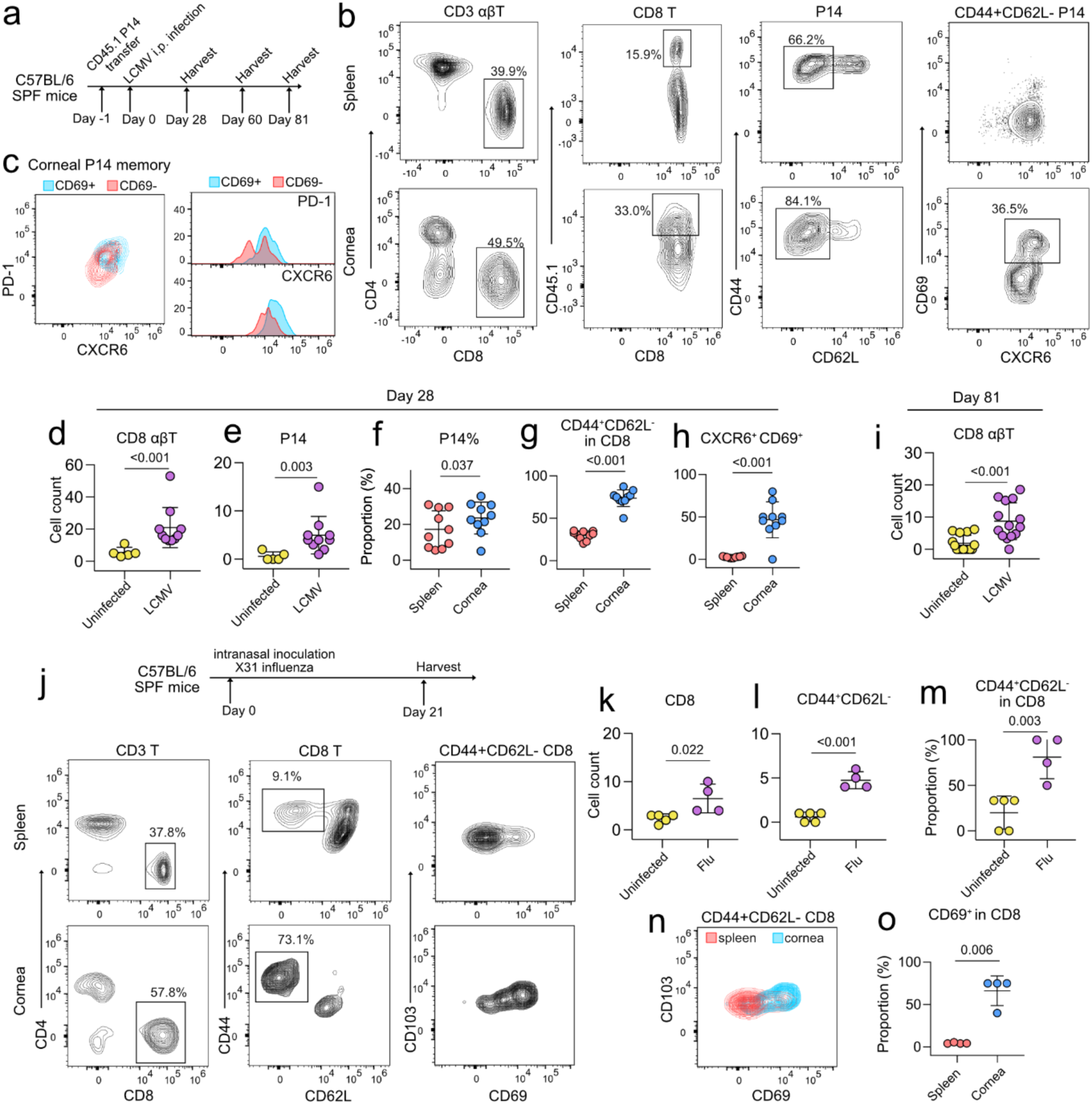
Systemic and respiratory viral infections induce the formation of corneal T_RM_ cells. (a) Experimental design involving mice adoptively transferred with CD45.1 or GFP P14 cells before LCMV infection prior to harvest of corneas and spleen at indicated timepoints. (b) Flow cytometry plots concatenated from 10 mice at day 28 after systemic LCMV infection. (c) At day 28 post-infection, CD69^+^ P14 cells in the cornea showed a shift toward a CXCR6⁺PD-1⁺ phenotype compared with CD69⁻ P14 cells. (d-e) Quantitative analysis showing more CD8^+^ T cells and P14 cells in the cornea at day 28 after systemic LCMV infection. and (f-h), a higher proportion of P14 cells memory and resident T cell phenotypes in the cornea compared to spleen. (i) Flow cytometry data reveals more CD8^+^ T cells in the cornea at day 81 after systemic LCMV infection compared to uninfected mice. (j) Experimental design of influenza infection mouse model where mice are infected with X31 intranasally and tissues harvested and analyzed via flow cytometry at day 21. (k-m) More CD8^+^ and CD44^+^CD62L^-^ T cells in the cornea at day 21 after X31 influenza infection compared to uninfected animals. (n-o) A higher proportion of effector-memory T cells co-expressing CD69 in the cornea compared to spleen. Statistical tests: Mann-Whitney test (d, e, i), paired *t* test (f, g, h and o), unpaired *t* test (k-m). Numbers above graphs indicate P values. Sample numbers: 5 (d-e, Uninfected; k-m, Uninfected), 10 (d-e, LCMV; f-h), 15 (i, Uninfected), 14 (i, LCMV), 4 (k-m, Flu; o). Data are shown as mean ± SD.

To further test our hypothesis, we employed a mouse model involving intranasal inoculation of influenza infection (X31 strain), which induces mild-to-moderate, non-lethal illness and recovery within two weeks (Fig. 4j). At 21 days post-influenza infection, there were significantly more CD8⁺ and CD44^+^CD62L^-^ T cells in the cornea compared to naïve animals (Fig. 4k-m); most of these corneal CD44^+^CD62L^-^ T cells also expressed CD69 (Fig. 4n-o). These findings lend additional support to the notion that systemic immune challenges can promote the establishment of T_RM_ cells in the cornea from populations of circulating effector cells.

### CXCR6 mediates non-redundant corneal T cell recruitment and establishment of corneal T_RM_

Since CXCR6 and CXCR3 have been shown to promote the recruitment, localization, and persistence of CD8^+^ T cells within non-lymphoid tissues^20, 24^, we hypothesized that these receptors may similarly facilitate T-cell entry into the cornea. Here, we used a CRISPR-Cas9 system to knock out *Cxcr6* or *Cxcr3* on P14 transgenic T cells to assess their role in T_RM_ formation in the cornea. We co-transferred wild-type (*sgCd19*) and CRISPR-knockout (*sgCxcr6* or *sgCxcr3*) P14 T cells. At eight days after systemic LCMV infection, we observed significantly less recruitment of *sgCxcr6* P14 cells to the cornea, relative to the spleen (Fig. 5a-b, Supplemental Fig. 4a). The *sgCxcr6* P14 cell population also showed a lower proportion of early T_RM_-phenotype cells (CD69^+^CD103^+^) compared with their *sgCd19* counterparts (Fig. 5c). In contrast, CXCR3 deficiency using either CRISPR-mediated knockout or genetic deficiency did not impair T cell entry into the cornea (Fig. 5d-e, Supplemental Fig. 4b-d). These findings identify CXCR6 as a modulator of T cell recruitment into the cornea after systemic immune activation.

**Figure 5.**
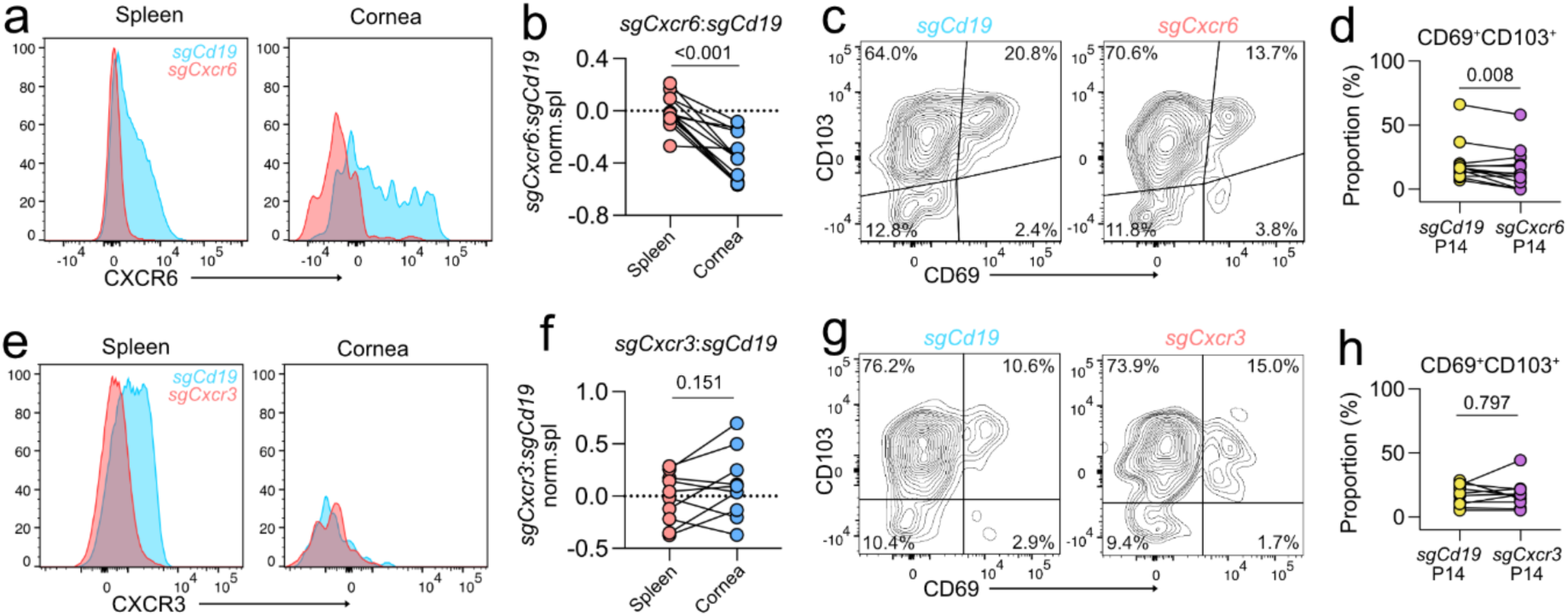
Corneal T_RM_ formation is dependent on CXCR6. CRISPR/Cas9 depletion of *Cxcr6* (a-d) or *Cxcr3* (e-h) in naïve P14 T cells. Control (*sgCd19*) and *sgCxcr6* or *sgCxcr3* treated P14 T cells were co-transferred into mice prior to LCMV infection and tissues harvested on day 8. (a) Histograms show CXCR6 expression in *sgCd19* and *sgCxcr6* P14 T cells. (b) Numbers of *sgCxcr6* and *sgCd19* treated P14 cells in the cornea relative to spleen. (c) Flow cytometry plots showing representative gating for CD103^+^CD69^+^ P14 T cells. (d) Frequency of CD69^+^CD103^+^ *sgCxcr6* P14 T cells compared to *sgCd19* P14 T cells. (e) Histograms show CXCR3 expression in *sgCd19* and *sgCxcr6* P14 T cells. (f) Numbers of *sgCxcr3* and *sgCd19* treated P14 cells in the cornea relative to spleen. (g) Flow cytometry plots showing representative gating for CD103^+^CD69^+^ P14 T cells. (h) Frequency of CD69^+^CD103^+^ *sgCxcr3* P14 T cells compared to *sgCd19* P14 T cells. Numbers above graphs (b, d, f, h) indicate P values from paired *t* tests. n = 8 (a), 13 (b-d), 5 (e), 10 (f-h).

## Discussion

For decades, the cornea has been considered a prototypical immune privileged tissue, owing to its lack of blood and lymphatic vessels and the perceived lack of surveillance by lymphocytes under steady-state conditions^1, 25^. Here, we show that the human cornea houses CD8^+^ T_RM_-like cells, at least some of which we propose are progressively primed by systemic immune challenge across the lifespan and accumulate in the cornea. At least in part, this process is governed by CXCR6-dependent recruitment or retention of T cells in the corneal tissue.

The identification of T_RM_ cells in barrier tissues has revolutionized understanding of regional tissue immunity^26^. CD8^+^ T_RM_ cell (CD69^+^CD103^+/-^) populations frequently develop following local infectious challenges, including in the skin, gastrointestinal tract and lung^27, 28, 29^. While mechanisms regulating T_RM_ cell formation in peripheral tissues are still being elucidated^30^, animal models show that systemic immune challenge can induce effector CD8 T cells to migrate to multiple non-lymphoid tissues and differentiate into T_RM_ cell phenotypes^13, 14, 31^. These findings suggested that T_RM_ cells can form in non-lymphoid tissues independently of local antigen recognition, which was further supported by T_RM_ cell formation in the skin following direct injection of *in vitro*-activated CD8^+^ T cells^32^. Here, we reveal that the cornea harbors a T_RM_ cell compartment that is shaped by prior systemic immune exposures. This concept aligns with recent evidence in other immune-privileged sites, such as the brain and the meninges, where T cells have been shown to share a consistent T_RM_ phenotype across donors with or without neurological diseases^33^.

While immune cell abundance in post-mortem tissues can be affected by a range of factors (e.g., tissue preservation time and digestion strategy, Supplemental Fig. 1g-h), our human intravital Fun-IVCM imaging data provide inferential *in vivo* support of this systemic immune-priming model. Time-lapsed imaging enables the identification of motile T cells in the living human corneal epithelium, which are identifiable by their distinct morphodynamic characteristics^5^. Based on the present study’s findings, we infer that T_RM_ cell populations represent a substantial proportion of these cells visualized in the human cornea. The positive correlation between age and corneal T cell density implies that the immune landscape of the steady-state human cornea undergoes progressive age-associated expansion. The progressive accumulation of T cells in the human cornea may also contribute to the phenomenon of ‘inflammaging’ at the ocular surface, whereby cumulative recruitment of resident T cells may increase the likelihood of chronic inflammatory conditions, such as dry eye disease, which are known to increase in prevalence with age^34^. Age-associated T_RM_ cell populations have been identified in the central nervous system in specific pathogen free laboratory mice, indicating that the T_RM_ cells may form by mechanisms other than infection, including neuroinflammation and age-related chemokine changes^35^. Surprisingly, negative correlations between biological age and CD103^+^ T cells have been reported in the endocervix and ectocervix, but not endometrium, in postmenopausal women^36^, suggesting the effect(s) of aging are tissue-dependent. Notably, age-associated increases in *in vivo* corneal intraepithelial T cell density within our human cohort were not evident in women. Supporting the notion that biological sex can differentially modify immune features in peripheral tissues, a recent study revealed that adult female mice have more resident T cells in their skin than males; while this sex-related difference was not observed in the small intestine^37^. These findings identify that the influence of sex-dependent factors on the expansion of T_RM_ cell populations in barrier tissues can be tissue-specific, with our data supporting the presence of such sex-driven differences in the human cornea. Further work is required to determine the effect of biological sex on corneal T_RM_ cell establishment, maintenance and function, and any associated implications for sex-dependent differences in corneal defense over the lifespan.

In the present study, murine pathogen infections provided a mechanistic framework for investigating how T_RM_ cells can develop in the cornea. Corneal T_RM_ populations (CD69^+^CD103^+^CXCR6^+^) were established, and persisted, in the absence of corneal pathology, following acute systemic and respiratory viral infection, indicating that the ocular surface microenvironment provides the necessary signals to support long-term cell residency and maintenance^19^. Importantly, we identify CXCR6 as a key chemokine receptor that is required for T cell entry into the cornea. This is consistent with work implicating CXCR6-CXCL16 interactions in T_RM_ cell positioning and retention in other non-lymphoid tissues^20, 21^, and with our observation that human corneal T cells express T_RM_-associated markers, including CXCR6. Another receptor, CXCR3, is often considered important for efficient T_RM_ formation in inflamed tissues, including skin and lung^24, 38^. CXCR3-CXCL10 has also been shown to have a critical role in generating the T_RM_ response to corneal HSV re-activation^39^. In contrast, we find that induction of T_RM_ cells in the cornea, as induced by systemic immune activation, is independent of CXCR3. These findings highlight that T_RM_ cell formation in the steady-state cornea is likely modulated by distinct chemokine axes that feature CXCR6-dependent mechanisms.

It is possible that corneal T_RM_ cells provide beneficial bystander protection to the ocular surface. In other barrier tissues, such as the skin, T_RM_ cells can rapidly produce cytokines and induce local immune responses, resulting in broad regional protection beyond antigen-specific cytotoxicity^31^. However, corneal T_RM_ cell populations may also contribute to chronic inflammation in ocular surface disorders or influence the outcome of ocular surface surgery, as recent studies have shown that persistent or aberrant T_RM_ cell activation can sustain local inflammatory cytokine production and amplify tissue immune responses independent of ongoing infection^40, 41^. Understanding the antigenic specificities, functional programs and clonal relationships of corneal T_RM_ cells will be essential for harnessing any protective potential while limiting immunopathology.

We acknowledge some limitations to the current work. While our findings establish the presence of resident T cell populations in the human cornea, further studies are needed to determine the clonal relationships and antigen specificities of these cells. There also remains the opportunity to interrogate the effector functions of corneal T_RM_ cells, including how they might contribute to protection and/or pathology in relevant disease models.

In summary, we show that the cornea is not an immunologically isolated tissue compartment, and is instead shaped by immune experience across the lifespan. By linking clinical *in vivo* imaging-based observational data with a mechanistic mouse model, our work provides a framework for understanding how tissue-resident T cells contribute to corneal health and identifies them as a potential target for therapeutic modulation of corneal immunity.

## Methods

### Human donor corneal tissues

Use of human corneal tissue for this study was approved by the University of Melbourne Human Research Ethics Committee (HREC, ID #24637). Corneal tissues were obtained from deceased individuals who donated their tissue for research use, from the Lions Eye Donation Service – CERA Biobank, Centre for Eye Research Australia, Royal Victorian Eye and Ear Hospital, University of Melbourne, or from the Eye Bank of South Australia Flinders Medical Centre, Southern Adelaide Local Health Network. Corneas were retrieved at 4 to 24 hours post-mortem and examined using a slit lamp biomicroscope at the Biobank, to confirm an absence of corneal scarring or neovascularization. Tissues were then stored in organ culture media for 2 hours to 30 days, prior to further processing. Donors with a history of ocular infection or trauma, or of major systemic infection, were excluded.

### Animals

All animal experiments were approved by The University of Melbourne Animal Ethics Committee (ID #25292). C57BL/6, P14 Ly5.1^Cg/Cg^, P14 GFP Ly5.1^Cg/Cg^, and P14 GFP Ly5.1^Cg/+^mice were bred at the Bioresource Facility of the Department of Microbiology and Immunology at The University of Melbourne. P14 GFP Ly5.1^Cg/+^ CXCR3^-/-^ mice were bred at the Walter and Eliza Hall Institute. Female mice were used at 6 to 12 weeks of age, except for the LCMV Day 81 post-infection experiment, where both male and female mice were used. P14 mice express a transgenic T cell receptor that recognizes the gp_33–41_ epitope of LCMV.

### Adoptive transfer of mouse P14 cells

P14 cell suspensions were acquired by mashing spleen and lymph nodes from P14 mice, through a 70-μm nylon cell strainer (BD Biosciences). Red blood cells were lysed with ammonium chloride-based buffers. C57BL/6 recipient mice were intravenously injected with 5 × 10^4^ P14 cells, at least 24 hours before infection. To determine whether CXCR3 is required for CD8^+^ T cell recruitment, CXCR3^-/-^ P14 cells were acquired from CXCR3^-/-^ P14 mice, followed by co-transfer of 1:1 ratio of CXCR3^-/-^ and CXCR3^+/+^ P14 cells, total of 5 × 10^4^ cells per mouse.

### CRISPR knockout

To determine whether CXCR3 or CXCR6 is required for CD8^+^ T cell recruitment, CRISPR– Cas9 editing of naive CD8^+^ T cells was performed, as described previously^42^. Single-guide RNAs (sgRNA) targeting: *Cxcr3* (5′-GAACAUCGGCUACAGCCAGG-3′, 5′- UGAGGGCUACACGUACCCGG-3′), *Cxcr6* (5′-UCUGUACGAUGGGCACUACG-3′, 5′- UGUGCCAAAGACCCACUCAU-3′) and *Cd19* (5′-AAUGUCUCAGACCAUAUGGG-3′, 5′-GAGAAGCUGGCUUGGUAUCG-3′) were purchased from Synthego (CRISPRevolution sgRNA EZ Kit). sgRNA/Cas9 RNPs were formed by incubating 0.3 nmol of each sgRNA with 0.6 μl Alt-R S.p. Cas9 nuclease V3 (10 mg/mL; Integrated DNA Technologies, cat#1081059) for 10 min at room temperature. Naive CD8^+^ T cells were negatively enriched from spleen of P14 mice by incubating cell suspensions with anti-CD4 (GK1.5), anti-CD11b (M1/70) anti- F4/80 (BM8), anti-Ter119 (TER-119) and anti-I-A/I-E (M5/114.15.2) monoclonal antibodies, followed by incubation with goat anti-rat IgG-coupled magnetic beads before removing bead-bound cells. A total of 1 × 10^7^ enriched T cells were resuspended in 20 μl of P3 (P3 Primary Cell 4D-Nucleofector X Kit; Lonza, cat#V4XP-3032), mixed with sgRNA/Cas9 RNP and electroporated using a Lonza 4D-Nucleofector system (DN100). Cells were rested for 30 min in a 96-well plate before direct transfer into recipient mice (co-transfer of 1:1 ratio of *sgCd19*:*sgCxcr3* or *sgCD19*:*sgCxcr6* CRISPR-edited naive P14 cells, total of 5 × 10^4^ cells per mouse). Adoptive transfer of CRISPR-edited P14 cells was performed at least seven days prior to infection.

### Mouse LCMV and influenza infection models

The Armstrong strain of LCMV was used. Mice were infected by intraperitoneal injection with 2 × 10^5^ plaque-forming units of LCMV Armstrong. For influenza infection, mice were infected intranasally with 10^4^ plaque-forming units of influenza A virus X31 (H3N2) in a volume of 30 μl. Mice were monitored and weighed daily for the first two weeks of the experiment, and then twice weekly. Corneal and/or spleen tissues were collected at Day 8, Day 28, Day 60, and Day 81 after LCMV infection, and at Day 21 after influenza infection.

### Corneal immunofluorescence staining and confocal imaging

Corneal tissues were fixed in Zambonis fixative at 4°C for 2-3h for immunofluorescence staining. Human corneas were radially segmented into six pieces to facilitate flat mounting. Corneal tissues were permeabilized in PBS containing 1% Triton X-100 for 1 hour, followed by blocking in PBS containing 2% BSA, 0.2% Triton X-100, and 1% Fc block for 1 hour at room temperature. Corneal tissues were then incubated overnight (4°C) in blocking buffer containing primary antibodies, followed by secondary antibody incubation for 2 hours at room temperature where required (antibodies used are listed in Supplementary Table S2). Stained tissues were flat mounted in low-fade mounting media. For human corneal tissues, a tile image of 1.39 to 2.46 mm^2^ area was acquired, for both the central and peripheral corneal regions, on a LSM980 confocal microscope (Carl Zeiss). For mouse tissues, one representative z-stack image (424x424 microns) of the central cornea and three images of the peripheral cornea were acquired on a LSM980 confocal microscope. Immune cell density quantification was performed by manual counting.

### Flow cytometry

Human donor corneas were dissected into central (<5mm diameter) and peripheral (5-9mm diameter) parts, with care to ensure that both the limbus and sclera were excluded (Supplemental Fig.1b). Mouse corneas were dissected using a 2.5mm trephine, to also exclude the limbus, and two corneas were pooled for each animal. Corneal tissues were minced into small pieces and digested using Liberase TL (0.25mg/mL, Roche) and DNase I (0.01mg/ml, Sigma Aldrich) (for human cornea), or Collagenase D (1mg/m, Sigma Aldrich l) and DNase (0.1mg/ml) (for mouse cornea) for 30 minutes at 37°C. The resulting cell suspension was collected, and fresh digestion enzymes were added every 10 minutes until no solid tissue remained. Mouse spleens were mechanically dissociated through a 70-μm nylon mesh. The single-cell suspensions were then stained with a panel of antibodies (Supplementary Table S2) for 30 minutes at 4°C, with single staining references prepared using compensation beads. Live cells were discriminated with a Zombie NIR or Zombie Aqua Fixable Viability Kit (Biolegend). Samples were acquired using Cytek Aurora (Cytek Biosciences), and FlowJo software (Tree Star) was used for analysis.

### Human participants

For the functional *in vivo* confocal microscopy (Fun-IVCM) *in vivo* imaging data, a pooled, retrospective analysis was performed on healthy (control) participant data from multiple studies involving participants seen at the University of Melbourne (UoM), between 2022 and 2026. Human research ethics committee (HREC) approvals were prospectively granted for each study (UoM HREC IDs #22828, #23535, #27067, #29215, #32768, #137830; Alfred Hospital HREC ID #191-17). All participants provided written informed consent to participate. The studies were conducted in accordance with the Declaration of Helsinki. The studies were originally designed to investigate ocular or systemic conditions, with each including a healthy (control) arm as a comparator. Some of these study populations have been reported previously^15, 43, 44^, whereas others are currently unpublished.

For the present work, analyses involved all individuals designated as healthy controls in the original studies that met the following inclusion criteria: aged ≥ 18 years, best-corrected visual acuity within normative ranges, and an absence of prior or current clinically significant ocular pathology. Exclusion criteria included a history of keratitis, ocular surgery, an ocular or systemic condition(s) that may affect corneal health (e.g., dry eye disease, contact lens wear, diabetes, auto-immune diseases), and a COVID-19 or influenza vaccination within the week prior to the study visit. Participant demographic information, including age and biological sex, was extracted from the study records.

Fun-IVCM time-lapsed videos were acquired, as described previously ^45^. In brief, topical anesthesia (0.4% oxybuprocaine hydrochloride; Bausch & Lomb) was applied to participants’ right eyes for intravital imaging using the Heidelberg Retinal Tomograph-III with Rostock Corneal Module (Heidelberg Engineering). Two representative corneal regions were imaged, comprising the whorl (inferonasal, paracentral cornea) and peripheral (∼2mm inferior to the whorl) cornea. For each region, repeated scans were acquired every 5.0 ± 1.5 minute, for 5 to 6 timepoints. Time-lapsed videos were registered using MATLAB (The MathWorks, version R2023a) using custom scripts ^45^. Corneal immune cell density was quantified by manual counting. Immune cell dynamic behaviors were measured using manual tracking in ImageJ for T cell mean instantaneous speed (by averaging instantaneous speeds across all time intervals) and DC dSEARCH index (derived by calculating the cumulative distance of dendrite extension and retraction of all dendrite tips from a single cell, normalized by time, then averaged across all time-intervals).

### Statistical analysis

For human donor tissue data, paired *t* test or Wilcoxon test was used to compare the central and peripheral corneal regions. For human *in vivo* imaging, as cell density data were right-skewed, a log10 transform was applied. Associations with age, sex and tissue region were assessed using linear mixed-effects models (participant ID as random intercept), with best fitting model selected by likelihood ratio tests. Correlations between participant age and immune cell densities were assessed separately for the whorl and peripheral regions using Pearson or Spearman correlation analysis based on the data normality. Comparisons between participants’ age sub-groups (younger: <40 years and older: ≥40 years) for human corneal immune cell densities were performed using Welch’s *t* test. Linear mixed-effects models were fitted to *in vivo* immune cell dynamic data at the participant level (with cell level data averaged per participant) and cell level. Model-predicted associations between participant age and immune cell dynamic features were also analyzed. Clinical data analyses were performed in R (version 4.5.2). For animal studies, unpaired *t* test or Mann-Whitney test was used for comparisons between groups, and paired *t* tests were used for comparisons between the cornea and spleen, using GraphPad Prism (version 11.0.0). A P value of less than 0.05 was considered statistically significant.

## Acknowledgements

We acknowledge the Lions Eye Donation Service – CERA Biobank, Centre for Eye Research Australia, Royal Victorian Eye and Ear Hospital, University of Melbourne. We acknowledge the Eye Bank of South Australia Flinders Medical Centre, Southern Adelaide Local Health Network. We thank Ms Simone Bituin, Dr Senuri Karunaratne, Ms Anna Lee, Dr Rajni Rajan, Ms Hanqing (Heidi) Wang and Ms Xinyuan Zhang for performing the Fun-IVCM imaging and clinical data collection, and Prof Andrew Metha and Dr Phillip Bedggood for the MATLAB scripts for clinical image registration. We thank Prof Linda Wakim for providing the influenza virus. We thank the Bioresources Facility and the Melbourne Cytometry Platform at the Peter Doherty Institute for technical support. Confocal imaging was performed at the Biological Optical Microscopy Platform (BOMP) Facility at The University of Melbourne.

This research was primarily funded by an ARC Discovery Project Grant awarded to LED, SNM and HRC (DP230102105). We also report funding from NHMRC Investigator Grants (2033906 to LED and 2017220 to SNM) and a University of Melbourne (UoM) McKenzie Fellowship to MW.

## Author contributions

MW: data curation, methodology, formal analysis, software, visualization, and writing—original draft, review, and editing. SCM: data curation, methodology, and writing—original draft, review, and editing. MHY: data curation, methodology, and writing—review and editing. KS: data curation, methodology, and writing—review and editing. SD: methodology and writing—review and editing. YOA: methodology. CYW: data curation and writing—review and editing. TNB: methodology. JRG: methodology and resources. BNN: data curation and writing—review and editing. LKM: methodology and resources. HRC: conceptualization, methodology, funding acquisition, supervision, and writing—original draft, review, and editing. SNM: conceptualization, methodology, funding acquisition, supervision, resources, and writing— original draft, review, and editing. LED: conceptualization, methodology, funding acquisition, supervision, resources, and writing—original draft, review, and editing.

## Disclosure

MW, HRC, SNM and LED are listed inventors on a patent relating to the clinical imaging method, owned by the University of Melbourne.

