## Supplemental materials for "Human cornea harbors tissue-resident memory T cells shaped by systemic immune activation, age and biological sex"

**Supplemental Information**

**Supplemental Table 1** Human donor information

| Age | Sex | Time interval (death to retrieval) | Time interval (death to tissue preservation) | General medical history | Ocular history |
|---|---|---|---|---|---|
| 73 | M | 24 h | 1 d | Parkinson's disease | NA |
| 63 | F | 12 h | 7 d | Glioblastoma | NA |
| 26 | F | 40 h | 30 d | Gunshot | Canthotomies - bilateral |
| 69 | M | 20 h | 20 d | COPD, cardiorespiratory event | NA |
| 61 | M | 17 h | 2 d | Renal cell carcinoma, hepatitis B, oral herpes infection | NA |
| 50 | M | 18 h | 4 d | Colorectal adenocarcinoma | NA |
| 72 | F | 7 h | 6 d | Type 2 respiratory failure, hypertension, hypothyroidism, obstructive sleep apnea, atrial fibrillation | Macular degeneration |
| 60 | M | 23 h | 5 d | No significant history | NA |
| 78 | M | 21 h | 4 d | Parkinson's disease, subarachnoid hemorrhage, previous stroke, chronic subdural hematoma, haemochromatosis, atrial fibrillation | NA |
| 76 | F | 17 h | 1 d | T2DM, hypertension, chronic kidney disease, heart failure | NA |
| 82 | F | 20 h | 2 d | Breast cancer, cecal adenocarcinoma, hypertension | Cataract surgery |
| 69 | F | 6 h | 1 d | T2DM, hypercholesterolemia, nausea, GORD, gastritis, osteoarthritis, pancytopenia | NA |
| 86 | F | 1 h | 1 d | Alzheimer’s disease | NA |
| 29 | M | 18 h | 4 d | Cerebral palsy, childhood hyaline membrane disease, focal right-sided epilepsy | NA |
| 63 | M | 18 h | 3 d | Coronary artery graft surgery, squamous cell carcinoma of tongue, hypertension | Fuchs endothelium dystrophy |
| 58 | F | 15 h | 1 d | Grand mal epilepsy, rhinitis medicamentosa, shingles, hepatitis C (negative RNA 2015), asthma, GORD, hyperprolactinemia, | NA |

| | | | | | |
|---|---|---|---|---|---|
| | | | | pulmonary embolism, polycystic ovary syndrome, irritable bowel syndrome | |
| 67 | F | 22 h | 8 d | Gastric cancer, colonic polyposis, GORD, hiatus, hypercholesterolemia, sensory neuropathy | NA |
| 55 | M | 14 h | 1 d | Cardiac arrest, knee problems managed with analgesics | NA |
| 77 | M | 29 h | 2 d | Parkinson’s disease, aortic valve replacement, permanent pacemaker | NA |
| 82 | F | 2 h | 1 d | Alzheimer's, congestive heart disease, hypertension diverticulosis | NA |
| 68 | M | 18 h | 5 d | VAD, T2DM, motor neuron disease | Phakic |
| 31 | M | 20 h | 2 d | Asthma, intravenous drug use | NA |
| 56 | F | 17 h | 10 d | Carotid artery dissection, COPD, hypertension | NA |
| 69 | M | 20 h | 2 d | VAD, asthma, respiratory failure, hyperlipidemia, solar ketosis, motor neuron disease | NA |
| 72 | M | 20 h | 7 d | VAD, motor neuron disease | NA |
| 53 | M | 35 h | 21 d | Intracranial cerebral hemorrhage, hypertension, T1DM, hepatic steatosis, chronic kidney disease | NA |
| 79 | F | 18 h | 18 d | Hypertension, hypercholesterolemia, T2DM, breast cancer | Cataract surgery |

Abbreviations: COPD, Chronic obstructive pulmonary disease; GORD, Gastro-esophageal reflux disease; T1DM, Type 1 diabetes mellitus; T2DM, Type 2 diabetes mellitus; VAD, Voluntary assisted dying.

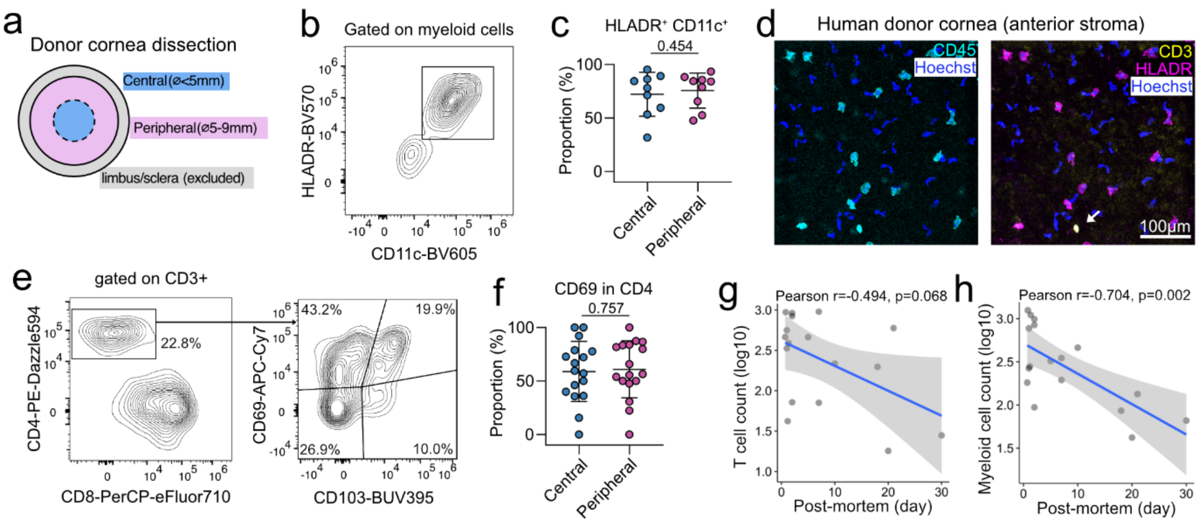


**Supplemental Figure 1. Immune cell populations in the healthy human cornea.** (a) Diagram showing the definition of central and peripheral human corneal regions for flow cytometry analysis. (b-c) Flow cytometry data concatenated from human donor corneas (n=9) reveals myeloid cells are predominantly HLA-DR$^+$CD11c$^+$. (d) Representative images from a 57-year-old female donor cornea show HLA-DR$^+$ myeloid cells dominating the immune compartment in the human corneal anterior stroma with occasional T cells observed (arrow). (e-f) Flow cytometry data concatenated from human donor corneas (n=17) demonstrates over half of CD4$^+$ T cells express CD69. (g) Correlation between human corneal T cell number and myeloid cell number (h) and tissue post-mortem time. Shaded areas (g and h) indicate 95% confidence intervals. Statistical tests: Paired *t* test (c and f), Pearson correlation test (g and h). Numbers above graphs indicate P values. Sample numbers: 9 (c) and 17 (f). Data (c and f) are shown as mean ± SD.

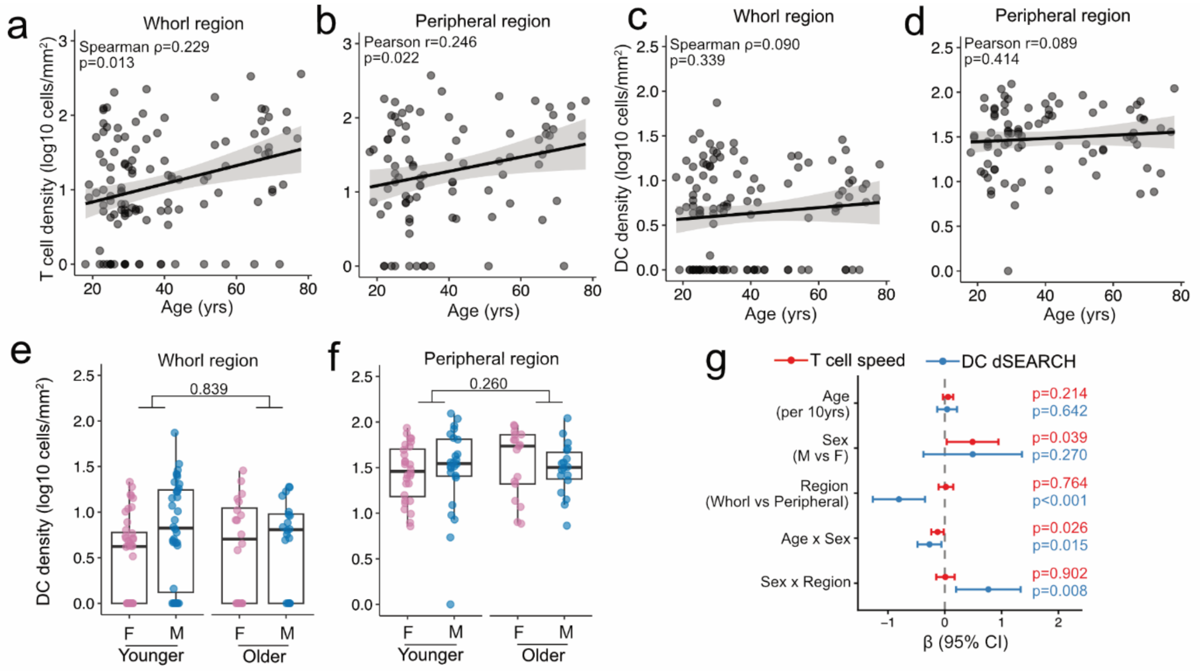


**Supplemental Figure 2. Human corneal T cell populations expand with age.** (a-b) Correlations between the density of putative human corneal intraepithelial T cells and participant age, in the whorl and peripheral regions. Shaded areas indicate 95% confidence intervals (CIs). (c-d) Correlations between the density of putative human corneal intraepithelial DCs and participant age, in the whorl and peripheral regions. Shaded areas indicate 95% CIs. (e-f) Human in vivo corneal DC density in younger (< 40 yrs) and older adults ($^{3}$ 40 yrs). (g) Forest plot demonstrating standardized β coefficients (with 95% CIs) from linear mixed-effects models fitted to cell-level dynamic data. Statistical tests: Spearman correlation test (a and c), Pearson correlation test (b and d), Welch's t test (e and f), linear mixed-effects model (g). Numbers above graphs indicate P values. Data (e and f) are shown as median with interquartile range.

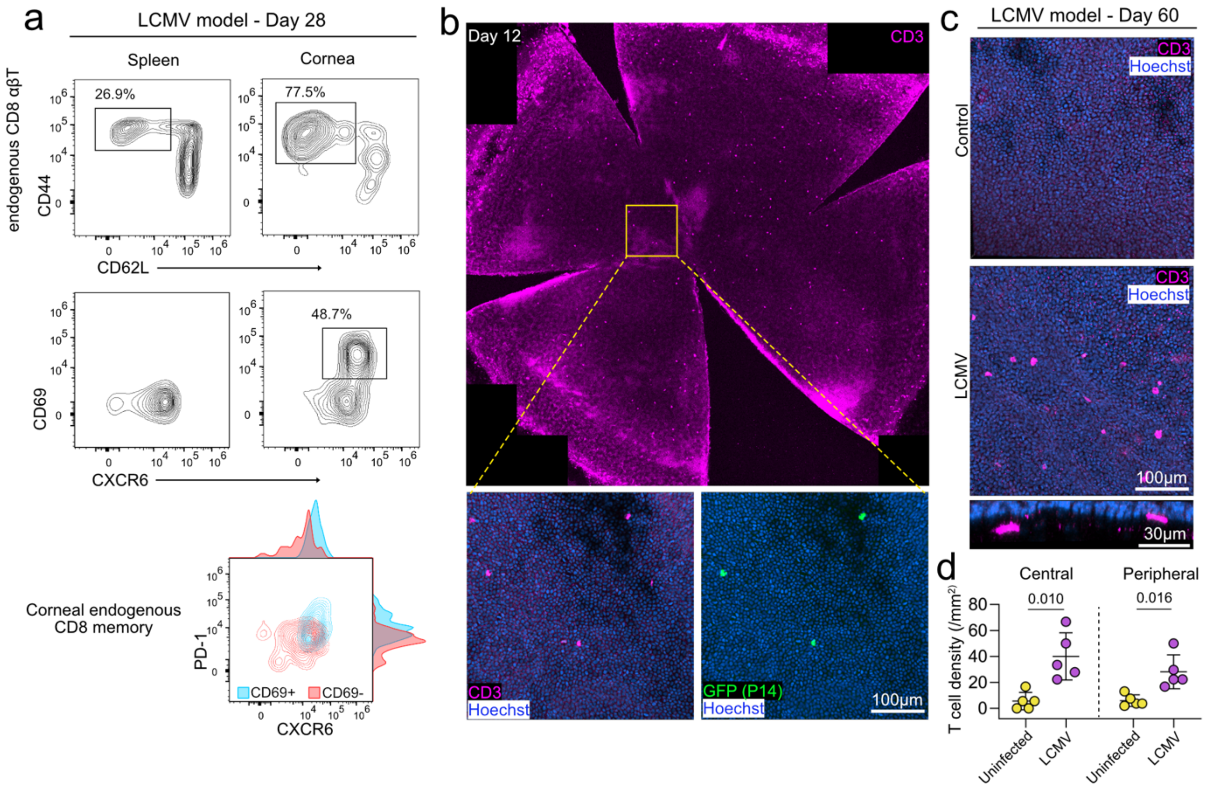


**Supplemental Figure 3. Systemic viral infection recruits T cells and generates T$_{RM}$ cells in the cornea.** (a) Flow cytometry data concatenated from 10 mice at day 28 after systemic LCMV infection where endogenous CD69$^{+}$ CD8$^{+}$ T cells in the cornea show a shift towards a CXCR6$^{+}$PD-1$^{+}$ phenotype compared with CD69$^{-}$ populations. (b) Representative corneal immunofluorescence images of a mouse with systemic LCMV infection at day 12, showing both P14 (GFP$^{+}$) and endogenous T cells in the cornea. (c) Representative mouse central corneal immunofluorescence images, showing more T cells in the cornea (present in both epithelium and anterior stroma) after systemic LCMV infection at day 60. (d) Quantitative data from uninfected and LCMV-infected mice at day 60, showing a higher T cell density in both the central and peripheral corneal regions. n = 5 mice. Numbers indicate P values from unpaired *t* test. Data are shown as mean ± SD.

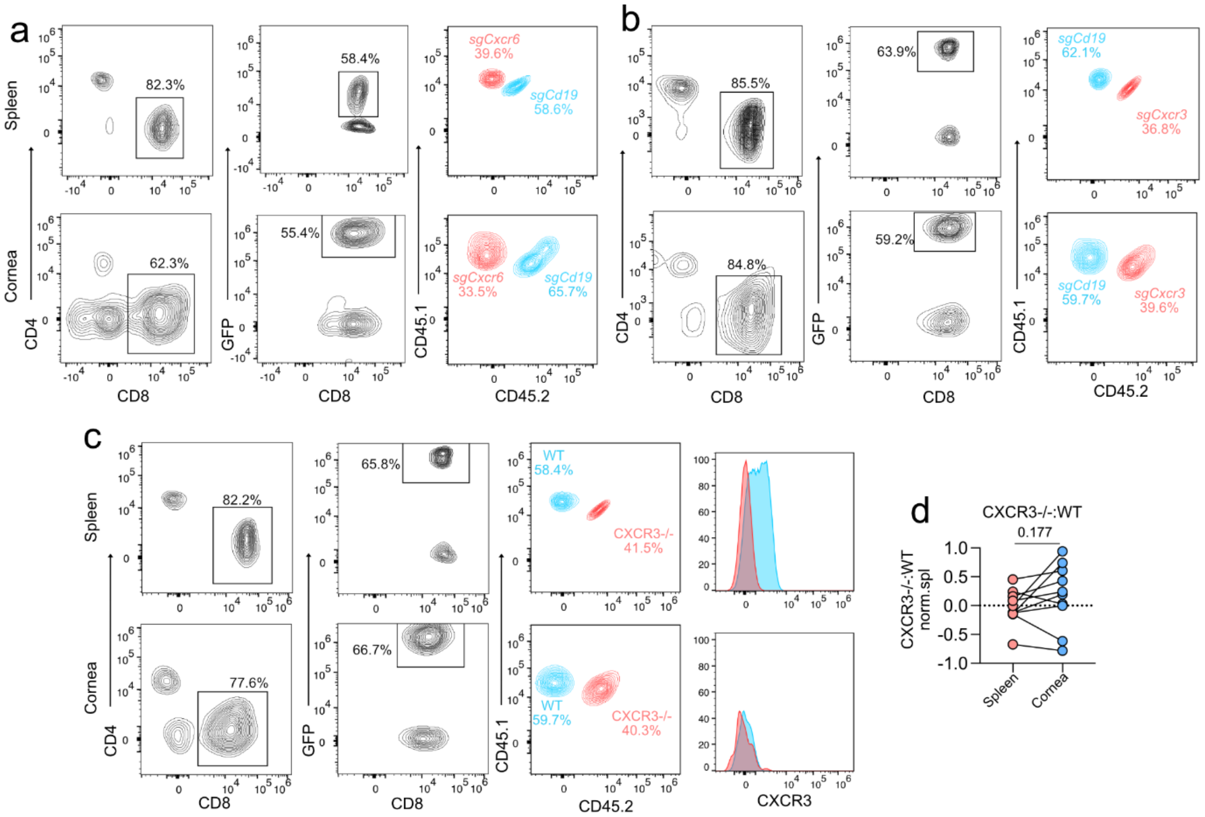


**Supplemental Figure 4. Corneal $T_{RM}$ populations form independently of CXCR3.** (a-b) Flow cytometry plots showing P14 T cell gating in a co-transfer model using wild-type (*sgCd19*) and CRISPR-knockout (*sgCxcr6 or sgCxcr3*) P14 T cells followed by systemic LCMV infection. Concatenated from 8 (a) or 5 (b) mice at day 8 after systemic LCMV infection. (c) Transfer of wild-type (WT) and $CXCR3^{-/-}$ P14 T cells followed by systemic LCMV infection. Flow cytometry plots concatenated from 5 animals at day 8 after systemic LCMV infection. (d) Ratio of $CXCR3^{-/-}$ and WT P14 cells in the cornea relative to spleen. n = 10 mice. Numbers above graphs indicate P value from paired t test.

**Supplementary Table S2** Antibodies used in this study.

| Antibodies | Source | Identifier |
|---|---|---|
| APC/Fire 750 anti-human CD45, clone HI30 | Biolegend | Cat# 304061 |
| Alexa Fluor 647 anti-human HLA-DR, clone LN3 | Biolegend | Cat# 327012 |
| Alexa Fluor 647 anti-human CD8, clone C8/144B | Biolegend | Cat# 372906 |
| Alexa Fluor 700 anti-human CD4, clone RPA-T4 | Biolegend | Cat# 300526 |
| Alexa Fluor 594 anti-human CD3, clone UCHT1 | Biolegend | Cat# 300446 |
| Alexa Fluor 488 anti-human CD69, clone FN50 | Biolegend | Cat# 310916 |
| Rabbit anti-human CD103, clone EPR4166(2) | Abcam | Cat# ab129202 |
| Cyanine3 (Cy3) goat anti-rabbit IgG, polyclonal | Invitrogen | Cat# A10520 |
| BUV395 anti-human CD103, clone Ber-ACT8 | BD Biosciences | Cat# 564346 |
| BUV496 anti-human CD20, clone 2H7 | BD Biosciences | Cat# 749954 |
| BUV737 anti-human CD3, clone UCHT1 | BD Biosciences | Cat# 612751 |
| BUV805 anti-human CD14, clone M5E2 | BD Biosciences | Cat# 612902 |
| BV421 anti-human CD45, clone HI30 | BD Biosciences | Cat# 563880 |
| BV570 anti-human CD45RA, clone HI100 | Biolegend | Cat# 304132 |
| BV605 anti-human CD11c, clone B ly6 | BD Biosciences | Cat# 563930 |
| BV711 anti-human CD56, clone HCD56 | Biolegend | Cat# 318336 |
| BV750 anti-human HLA-DR, clone L243 | Biolegend | Cat# 307672 |
| BV785 anti-human TCRab, clone IP26 | Biolegend | Cat# 306742 |
| FITC anti-human TCRgd, clone B1.1 | Invitrogen | Cat# 11-9959-42 |
| PerCP anti-human CD19, clone H1B19 | Biolegend | Cat# 302228 |
| PerCP-Cy5.5 anti-human CD45RO, clone UCHL1 | BD Biosciences | Cat# 560607 |
| PerCP-eFluor710 anti-human CD8a, clone SK1 | Invitrogen | Cat# 46-0087-41 |
| PE-Dazzle594 anti-human CD4, clone OKT4 | Biolegend | Cat# 317448 |
| APC anti-human CXCR6, clone K041E5 | Biolegend | Cat# 356006 |
| APC-cy7 anti-human CD69, clone FN50 | Biolegend | Cat# 310914 |
| BUV395 anti-mouse B220, clone RA3-6B2 | BD Biosciences | Cat# 563793 |
| BUV805 anti-mouse CD4, clone RM4-4 | BD Biosciences | Cat# 741913 |
| BV480 anti-mouse CD103, clone M290 | BD Biosciences | Cat# 566118 |
| BV570 anti-mouse Ly6C, clone HK1.4 | Biolegend | Cat# 128030 |
| BV650 anti-mouse TCRb, clone H57-597 | BD Biosciences | Cat# 742483 |
| BV711 anti-mouse PD-1, clone 29F.1A12 | Biolegend | Cat# 135231 |
| BV785 anti-mouse CD45.1, clone A20 | Biolegend | Cat# 110743 |
| FITC anti-mouse CXCR6, clone SA051D1 | Biolegend | Cat# 151108 |
| BUV737 anti-mouse CD44, clone IM7 | BD Biosciences | Cat# 612799 |
| PerCP anti-mouse CD45, clone 30-F11 | BD Biosciences | Cat# 557235 |
| PerCP-Cy5.5 anti-mouse CXCR3, clone CXCR3-173 | Biolegend | Cat# 126514 |
| PE anti-mouse CD3e, clone 145-2C11 | BD Biosciences | Cat# 553064 |
| PE-Dazzle594 anti-mouse CD45.2, clone 104 | Biolegend | Cat# 109846 |
| PE-Cy5 anti-mouse CD69, clone H1.2F3 | Biolegend | Cat# 104510 |
| PE-Cy7 anti-mouse CD62L, clone MEL-14 | Invitrogen | Cat# 25062182 |
| PE anti-mouse CXCR6, clone SA051D1 | Biolegend | Cat# 151104 |
| PE-Fire700 anti-mouse CD3, clone 17A2 | Biolegend | Cat# 100272 |
| Spark NIR 685 anti-mouse CD8a, clone 53-6.7 | Biolegend | Cat# 100782 |
| Alexa Fluor 647 anti-human CD3, clone KT3.1.1 | Biolegend | Cat# 155609 |
| Anti-mouse CD4, clone GK1.5 | Biolegend | Cat# 100401 |
| Anti-mouse CD11b, clone M1/70 | Biolegend | Cat# 101201 |
| Anti-mouse F4/80, clone BM8 | Biolegend | Cat# 123101 |
| Anti-mouse Ter119, clone TER-119 | Biolegend | Cat# 116201 |

| | | |
|---|---|---|
| Anti-mouse I-A/I-E, clone M5/114.15.2 | Biolegend | Cat# 107601 |
| Goat anti-rat IgG-coupled magnetic beads | Qiagen | Cat# 310107 |
